# Polymorphic SINEC_Cf Retrotransposons in the Genome of the Dog (*Canis familiaris*)

**DOI:** 10.1101/2020.10.27.358119

**Authors:** Sara E. Kalla, Hooman K. Moghadam, Max Tomlinson, Allison Seebald, Jeremy J. Allen, Jordan Whitney, Jessica D. Choi, Nathan B. Sutter

## Abstract

The dog is an exciting genetic system in which many simple and complex traits have now been mapped. For many traits the causal mutation is a polymorphic SINE. To investigate the genome-wide pattern of young SINEC_Cf insertions, we sampled 62 dogs representing 59 breeds and sequenced libraries enriched for SINE flanks. In each dog we detect an average of 10,423 polymorphic loci and all together the libraries identify 81,747 putative polymorphic SINEs. We validated 184 SINEs inserted in protein-coding exons, untranslated regions, introns and intergenic sequence. In dogs both SINEC_Cf and LINEs exhibit a strand bias in introns where antisense copies are more frequent. Antisense polymorphic SINEs also have a higher density in introns. Both SINEs and LINEs drop to very low density near exons. Both sense and antisense polymorphic SINEs also drop to low density upstream of coding exons but not downstream. Antisense polymorphic SINEC_Cfs upstream of coding exons are known to cause narcolepsy, merle, and progressive retinal atrophy in dogs. In other mammals SINE pairs in inverted orientation disrupt gene expression. We find inverted pairs of SINEC_Cf are rare in both introns and intergenic sequence when the two SINEs are separated by less than 100 bp. The lack of inverted pairs is even more pronounced when the SINEs have high sequence identity. Intronic and intergenic LINE pairs show similar patterns. Polymorphic SINEs rarely pair with either SINEC_Cf or SINEC_Cf2. Overall, the high insertion rate of SINEC_Cf provides a natural mutagenesis screen in the dog genome.

## Introduction

Dogs serve as a fantastic model system for many questions in genetics (Ostrander *et al.* 2000; Steenbeek *et al.* 2016). Most dogs in the world are feral or “village” dogs but millions of dogs are purebred, the products of intense selection within reproductively isolated lines. Pure breeds usually have bottlenecks at founding and often have been maintained as small populations throughout their development in the 19^th^ and 20^th^ centuries. Furthermore, many breeds are selected for extreme morphologic traits including small and large body size, body thickness variation, snout length, and limb length. There are also diverse coat colors, patterns, lengths, and curls selected within breeds. Unfortunately, most pure breeds also carry specific heritable disease risk factors.

Much progress has been made in identifying the genetic basis of both simple and complex traits in the dog (Boyko *et al.* 2010). The causal mutations for quite a number of traits turn out to be polymorphic retrotransposons. For example, a partial long interspersed element 1 (LINE1) copy interrupts an exon of *GRM1* in the coton de Tulear breed to cause neonatal ataxia (Zeng *et al.* 2011). Truncated polymorphic L1s in introns are reported to cause brachycephaly in many breeds (Marchant *et al.* 2017), muscular dystrophy in Pembroke Welsh corgis (Smith *et al.* 2011), and hemophilia B in German wirehaired pointers. In addition, there are reports of dog disease and other traits caused by polymorphic insertion of the short interspersed element, SINEC_Cf. This Can-SINE, first identified in the dog, is present in carnivores (Minnick *et al.* 1992; Coltman and Wright 1994; Das *et al.* 1998) and there are 171,386 copies in the dog reference genome.

A SINEC_Cf insertion present in herding breeds causes the merle coat pattern and associated eye and ear disorders. The SINE is inserted in an intron upstream of *PMEL* exon 11 at the exon/intron junction and disrupts splicing (Clark *et al.* 2006). The poly A track in the SINE’s tail is highly mutable and several distinct pigmentation alleles exist (Murphy *et al.* 2018). Narcolepsy in Doberman pinschers is also caused by a SINEC_Cf upstream of an exon; the SINE is 35 bp upstream of *HCRTR2* exon 4 (Lin *et al.* 1999). In both narcolepsy and merle the causative SINE is in an antisense orientation to the gene. This is also true for a polymorphic SINEC_Cf that causes progressive retinal atrophy in Tibetan spaniels and terriers. The SINE is inserted in antisense orientation 15 bp upstream of a coding exon in *FAM161A* and leads to exon skipping (Downs and Mellersh 2014). A polymorphic SINEC_Cf in an intron of *ASIP* that is one half of an inverted pair reportedly contributes to the black-and-tan coat pattern (Dreger and Schmutz 2011). In addition, SINEC_Cf insertions within coding exons cause retinal degeneration in Norwegian elkhounds (Goldstein *et al.* 2010), centronuclear myopathy in Labrador retrievers (Pelé *et al.* 2005), and a human Warburg-like syndrome in Alaskan huskies (Wiedmer *et al.* 2016). Thus, SINE disruption of genes is an important feature in the dog genome.

SINEs act similarly in other species. SINE disruption of human genes can occur in several different ways, for example via insertion within a coding exon, within untranslated sequence, and as part of an inverted pair (Cook *et al.* 2011; Taskesen *et al.* 2012; Anwar *et al.* 2017). The mouse genome is also altered by SINE activity (Ponicsan *et al.* 2010; Gagnier *et al.* 2019). In both genomes there is evidence for SINE “hazardous zones” (Zhang *et al.* 2011) defined by SINE exclusion and bias in frequency according to strand. In humans, *Alu* elements are involved in alternative splicing, RNA editing, and regulation of translation (Ponicsan *et al.* 2010; Daniel *et al.* 2015). In mice, a full-length B2 SINE insertion in the 5’ UTR of *ALAS1* causes a decrease in expression (Chernova *et al.* 2008), while a SINE insertion in the 3’ UTR of *COMT* adds a polyadenylation site, shortening the mRNA and increasing the transcription rate (Li *et al.* 2010). *Alus* are also responsible for gene silencing via double-stranded mRNA (dsRNA) formation in the 3’ UTR (Chen and Carmichael 2008). Furthermore, SINE elements can cause dsRNA formation and associated exon skipping; this occurs when two SINEs, such as *Alu* elements, are in opposite orientation (Lev-Maor *et al.* 2008), often called an “inverted repeat”.

The alterations in gene expression caused by SINEs can have significant effects on phenotypes and many disease-causing insertions have already been documented. In humans, retrotransposons cause numerous genetic diseases including cancer, Fukuyama muscular dystrophy, Dent’s disease, and Alström syndrome (Hancks and Kazazian 2016). The genome-scale analyses completed in human, mouse, and other species (Adelson *et al.* 2009; Ivancevic *et al.* 2016) will be rewarding to replicate in still more mammals so that we can better understand the evolution of genomes.

Wang and Kirkness addressed the question of SINE activity in the dog on a genome scale (Wang and Kirkness 2005) and identified more than 10,000 polymorphic SINEC_Cfs (which they termed “bimorphic”) despite the fact that only a few genomes were being compared. Combined with the evidence for polymorphism from trait mapping studies in dogs, these data argue that SINEC_Cf retrotransposition activity is very high and/or has been high in the recent past. With this in mind we endeavored to discover polymorphic SINEs in the dog genome to produce a dataset with which to assess the likely impact of SINEs on dog gene expression and traits. Toward this end we have generated and sequenced libraries of SINEC_Cf insertions in purebred dogs.

## Materials and Methods

### Ethics Statement and Sample Collection

Dogs were sampled with signed consent from owners under a protocol approved by the institutional animal care and use committee at Cornell University. Sampled dogs were judged to be purebred by owner statements on intake forms and by inspection of provided pedigrees. Genomic DNA was extracted by a Gentra-like protocol from whole blood that had been collected in EDTA or acid citrate anti-coagulant collection vials, shipped at ambient temperature and then stored for up to three weeks (but usually less than one week) at 4 °C. To lyse red blood cells, five volumes of ammonium chloride lysis solution (2 ml 0.5 M EDTA pH=8; 20 ml 0.5 M sodium bicarbonate; 8.3 g ammonium chloride; q.s. 1000 ml water and autoclave) was added to one volume of whole blood and mixed gently for 20 minutes. Samples were then centrifuged at 2000 x g for seven minutes, the supernatant was decanted from the small white pellet, and the pellet was resuspended in the residual liquid. One volume of white blood cell lysis solution (10 ml 1 M TRIS pH=8; 2 ml 0.5 M EDTA pH=8, 938 ml water; 50 ml 10% SDS added gently then filter sterilize) was added and samples were gently mixed overnight. On day two, 0.4 volumes of 10 M ammonium acetate solution were added and samples were vortexed at high speed to precipitate protein. Samples were sometimes placed on ice at this stage then they were centrifuged at 3300 x g for 10 minutes, the supernatant was poured off to a clean tube, one volume of 2-propanol was added to the supernatant, and the tube was gently mixed until DNA strands became visible and formed a defined white mass. DNA was pelleted by centrifugation at 2500 x g for five minutes and pellets were retained and washed with 1 ml of 70% ethanol followed by another centrifugation at 2500 x g for three minutes. The ethanol supernatant was then poured off and the DNA was air dried, resuspended in 10 mM tris at pH=8, and quantified by absorbance in a spectrophotometer. Sample, dog, pedigree, and breed records were stored in a database (Powell *et al.* 2010).

### SINE Library Production

Step 1 Extension. To produce libraries enriched for genomic sequences flanking SINEC_Cf insertions we first did second strand synthesis by hybridizing primer A855F (Table S1), which contains a 5’-biotin, onto each dog genomic DNA sample. Extension reactions were set up on ice and contained 10 μg of gDNA in a 50 μl reaction volume with 1x NH_4_ magnesium-free PCR buffer, 3 μl platinum taq DNA polymerase (Thermo Fisher Scientific), 200 μM dNTPs, 1.5 mM MgCl_2_, and primer A855F at a concentration of 200 nM. Extension reactions were split into 6.25 μl aliquots in each of eight wells and were overlaid with 8 μl of mineral oil. Reactions were manually cycled between two thermal cycler blocks set at 95 °C and 55 °C to ensure rapid temperature shifts and keep the extension times suitably short. After an initial incubation at 95 °C for two minutes, reactions were cycled 20 times at 55 °C for 10 seconds followed by 95 °C for one minute. The final incubation was 95 °C for two minutes. We calculated that the A855F primer would be in limiting supply in this reaction given the mass of gDNA in the reaction and the estimated number of loci at which the primer would bind. Thus relatively few of the biotinylated molecules in the reaction would be expected to be extension-less primers.

Step 2 Size Selection and Capture to Beads. All of the 50 μl extension reaction from a single sample was pooled, electrophoresed through a 1% TAE low melt agarose gel and size selected by cutting out the gel block containing the 160-300 bp size range. The DNA molecules were recovered by digesting the agarose gel block with one unit of agarase per 100 μg of agarose in a one hour incubation at 42 °C. Next, streptavidin-coated magnetic beads (Invitrogen, DYNAL) were used to capture the 5’-biotinylated extension products. Following the manufacturer’s guidelines, 10 μl of beads were washed three times in 100 μl of 1x bead wash buffer (10 mM Tris-HCl at pH=7.5, 1 mM EDTA and 2 M NaCl) then added to the agarase-digested gel block and incubated one hour at room temperature with shaking. Beads were captured in a magnetic field for five minutes then washed three times with 100 μl 1x bead wash buffer and rinsed with 100 μl PCR-grade water.

Step 3 Polyadenylation. Beads containing the ssDNA extension products were suspended in 10 μl of polyadenylation master mix containing 2 units of terminal transferase TdT (New England Biolabs), 1x TdT buffer, 0.25 mM CoCl_2_, and 100 μM dATP. Beads were incubated 90 minutes at 37 °C then washed three times with 1x bead wash buffer and rinsed with water, as above, before being suspended in 10 μl PCR-grade water.

Step 4 Poly-T Primed Second Strand Synthesis. All of the 10 μl polyadenylated beads were then used as template for a second strand synthesis reaction in 40 μl with 0.16 μl 5 U/ μl taq (Biolase), 1x Biolase KCl reaction buffer, 4.5 mM MgCl_2_, 800 μM dNTPs, and 1 μM poly-T primer A858F (Table S1). Primer A858F’s 3’-end nucleotides between positions 2-21 are all ‘T’. Because the polyadenylation reaction may add 100 or more ‘A’ nucleotides onto the 3’ ends of the bead-captured ssDNA, we designed A858F to have a non-‘T’ 3’-most nucleotide (any of ‘A’, ‘C’, and ‘G’) to anchor primer hybridization to the bead-proximal end of the poly-A run. After a hold at 94 °C for 30 seconds, the second strand synthesis reaction continued for 15 cycles at 94 °C for 30 seconds followed by primer annealing at 55 °C for 40 seconds and then extension at 72 °C for 1 minute. After at least one minute in a final hold at 94 °C the samples were removed from the cycler while hot and supernatant containing the synthesized second strands was recovered from the beads in a magnetic field.

Step 5 Introduce Barcodes and Illumina Forward Adaptor. One of the 16 barcoded primers, e.g. A1459F (Table S1), at a final reaction concentration of 1 μM was added to 10 μl of the above supernatant and seven cycles of PCR were carried out: hold 94 °C for 30 seconds then seven cycles at 94 °C for 30 seconds followed by 55 °C for 40 seconds and finally 72 °C for 1 minute. A final hold at 72 °C was five minutes long.

Step 6 PCR With Short Primers. Rounds of PCR with short oligonucleotides produce double stranded products and ensure they can be readily separated from the oligonucleotides. We added 10 μl of the Step 5 material into a 40 μl reaction containing 2 units of taq DNA polymerase (Biolase), 1x KCl reaction buffer (Biolase) and final reaction conditions of MgCl_2_ at 4.5 mM, dNTPs at 800 μM and primers A880F and A880R (Table S1) at 400 μM. After a hold at 94 °C for 30 seconds, the reaction was cycled 12-20 times at 94 °C for 30 seconds, 55 °C for 40 seconds and 72 °C for 90 seconds. A final extension at 72 °C was five minutes long. We cleaned the resulting PCR amplicons by following the manufacturer’s protocol with a PCR purification kit (Qiagen).

Step 7 Final Pippin Size Selection, Pooling and Sequencing. Small aliquots from each library were assessed for size distribution by agarose gel electrophoresis, then 16 libraries with distinct barcodes were pooled and size-selected by Pippin prior to loading into an Illumina HiSeq sequencer lane for single-end 100 bp sequencing with the standard Illumina adaptor primer. A total of four 16-plex lanes of sequence data were collected.

### SINE Library Parsing

After sequencing, we utilized a custom perl script to assign fastq sequence reads to the 16 samples in each of the four HiSeq lane datasets. The script checks for at most one mismatch to one of the barcodes for the 15 bases of each sequence read that start at the barcode and go 3’ (all barcode pairs have edit distances of three or higher). The script trims away the barcode and SINE hybridizing portion of the sequence read, as well as any poly-A sequence and Illumina adaptor sequence, if present, at the 3’ end of the read. The script produces a pair of sequences for each read: the “full” sequence contains 17 bases at the head end of the SINE plus the non-SINE flanking sequence. The “flank” sequence has the 17 SINE bases removed. Both the full and flank reads are independently aligned to the dog CanFam 3.1 reference genome (below). A total of 63 dog libraries were originally sequenced but a library from a Brussels griffon was represented in the pooled sequences at less than 1% the rate of other libraries and it was therefore removed from further analysis, leaving 62 libraries.

Before aligning our SINE sequence reads to the dog genome we quality filtered them in three ways. First, we checked for low complexity with PrinSeq, which removed on average approximately 20% of each sample’s sequence reads. Second, we repeat masked the non-SINE flanking portion of each read. Third, we have observed that many dog SINEC_Cf copies in the dog reference genome contain identical or nearly identical 17-mers at their head ends. We therefore gave each sequence read an integer score representing the number of SINEs in the dog reference genome that contained a 17-mer identical to the read. We removed from analysis all reads with 17-mers found in fewer than 1000 CanFam 3.1 SINEC_Cf copies.

Each flank and full sequence read was independently aligned to the dog CanFam 3.1 reference genome using the burrows wheeler aligner (BWA) with allowance for up to two mismatches in the alignment (Li and Durbin 2009). The resulting alignments were then filtered for flank reads that uniquely mapped to the reference genome. Parsed read sequences and alignment data were uploaded to a MySQL database that also contained the annotated SINEs and LINEs from the reference genome as well as the Ensembl 99 gene models for the dog. SINE alignments were assigned to an existing SINE record if they mapped on the same chromosome and strand and to within six base pairs of a SINE already in the dataset. This “fuzzy matching” is conservative with respect to the number of SINE insertion loci called and was used to efficiently match SINEs even when alignments were off by one or a few bp due to sequence errors or when the reference genome annotation misclassified the last few bases at the head ends of SINEs in the reference genome.

### SINE Analysis

Definition of Polymorphism. We defined polymorphic SINEs as those having evidence of insertion in one or more of our SINE libraries combined with a lack of any SINE annotation at that locus in the dog CanFam3 reference genome assembly.

Orientation Bias in Transcripts. We uploaded ensembl gene models from the UCSC genome browser into our sines database. We estimated promoters as extending 500 bp from the start of the 5’ UTR. To count SINEs within gene features we first collapsed all features such that each bp in the genome could be part of at most one gene feature. We kept feature annotations according to a hierarchy in which coding exons are always kept and coding exon > 5’UTR > 3’UTR > promoter > intron > intergenic. We counted the number of SINE insertions in each gene feature and tracked the relative orientation of the SINE and the gene. We also tallied counts for SINE inverted pair occurrences within gene features and intergenic sequence.

## Results

With an aim to understand the pattern of SINEC_Cf insertions in the dog genome, we collected Illumina HiSeq single-end 100 bp sequence reads from 62 dog DNA libraries. A total of 59 dog breeds are included in the dataset: 58 breeds are represented by a single dog and one breed, boxer, with four dogs. The libraries are enriched for sequences flanking the dog SINE retrotransposon sub-type SINEC_Cf and are generated by hybridizing a DNA primer (Table S1) to conserved sequence in the head end of SINEC_Cf (Figure 1).

**Figure 1.**
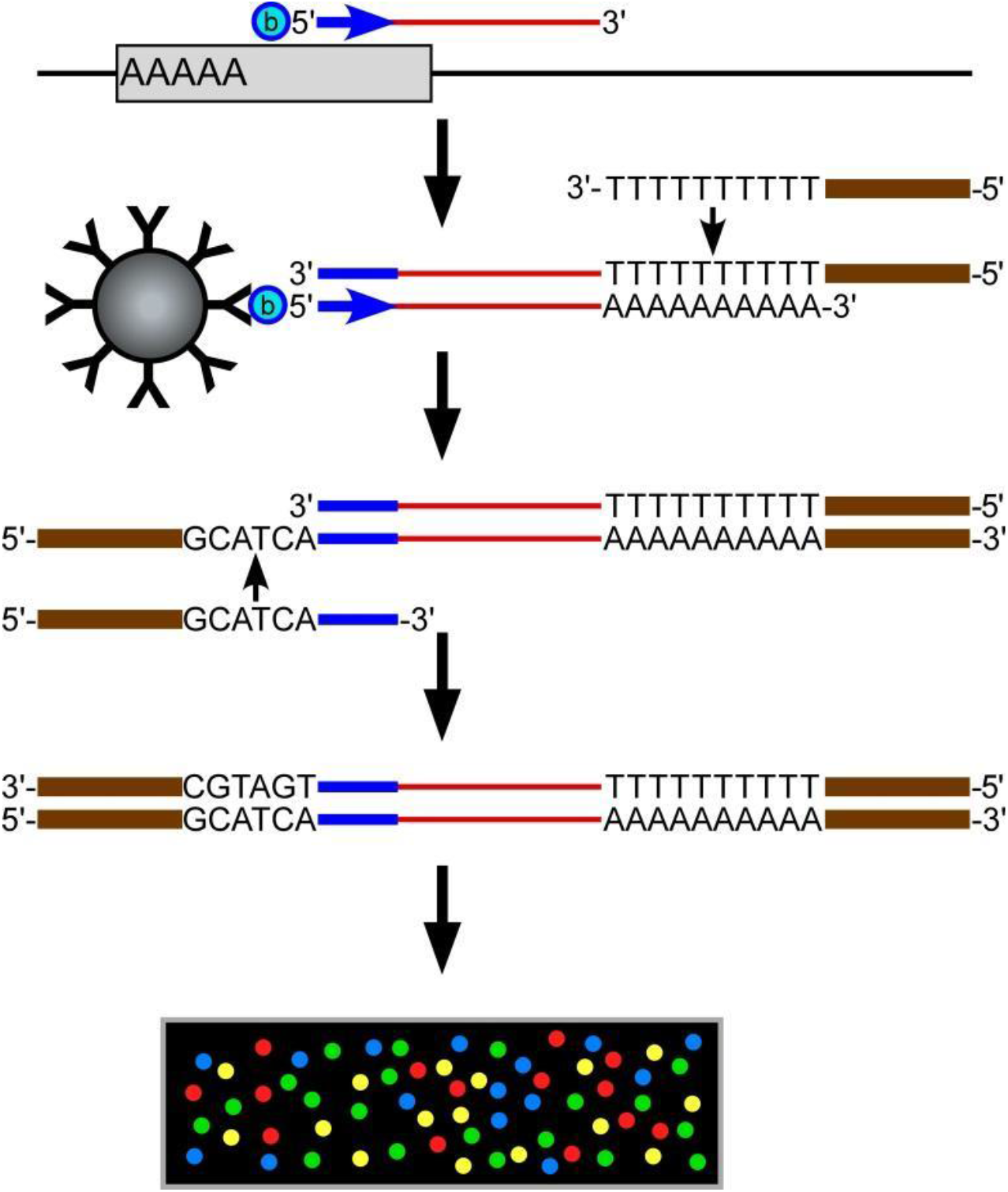
Construction of libraries enriched for sequences flanking the head end of SINEC_Cf copies. DNA synthesis is primed from dog genomic DNA with a biotinylated oligonucleotide that hybridizes to conserved sequence in SINEC_Cf. The primer’s 3’ end hybridizes to base 18 within the SINE. DNA extension products are captured to streptavidin-coated magnetic beads, polyadenylated, and used for 2^nd^ strand synthesis. After recovery from the beads, one of 16 barcoded primers tailed with Illumina HiSeq forward adaptor sequence is used for several rounds of PCR. After quality assessment by gel sizing, 16 libraries are pooled and subjected to one lane of 100 bp single-end sequencing.

After quality filtering, an average of 2.7 million sequence reads were collected from each library (minimum = 624,116, maximum = 4,938,963; Table 1). Libraries detected an average of 118,835 distinct loci (range: 95,156 in a bull terrier to 143,836 in a whippet), the majority of which are annotated SINEC_Cfs in the reference genome (Table 1). Each library detects about half of the 165,130 SINEC_Cfs annotated in the non-chrUn reference genome (minimum = 44.1% of reference SINEC_Cf loci, maximum = 58.8%; Figure 2). In aggregate our libraries detect 122,556 (74.2%) of all SINEC_Cfs in the reference genome. Many of the remaining reference SINEC_Cf loci probably can’t be found using this short read alignment strategy. We randomly chose 20 reference loci that our libraries failed to detect and found in 19 of them repeated sequences within 65 bps of the SINEC_Cf head end that would cause us to filter out any sequence reads emanating from the SINE.

**Table 1.**
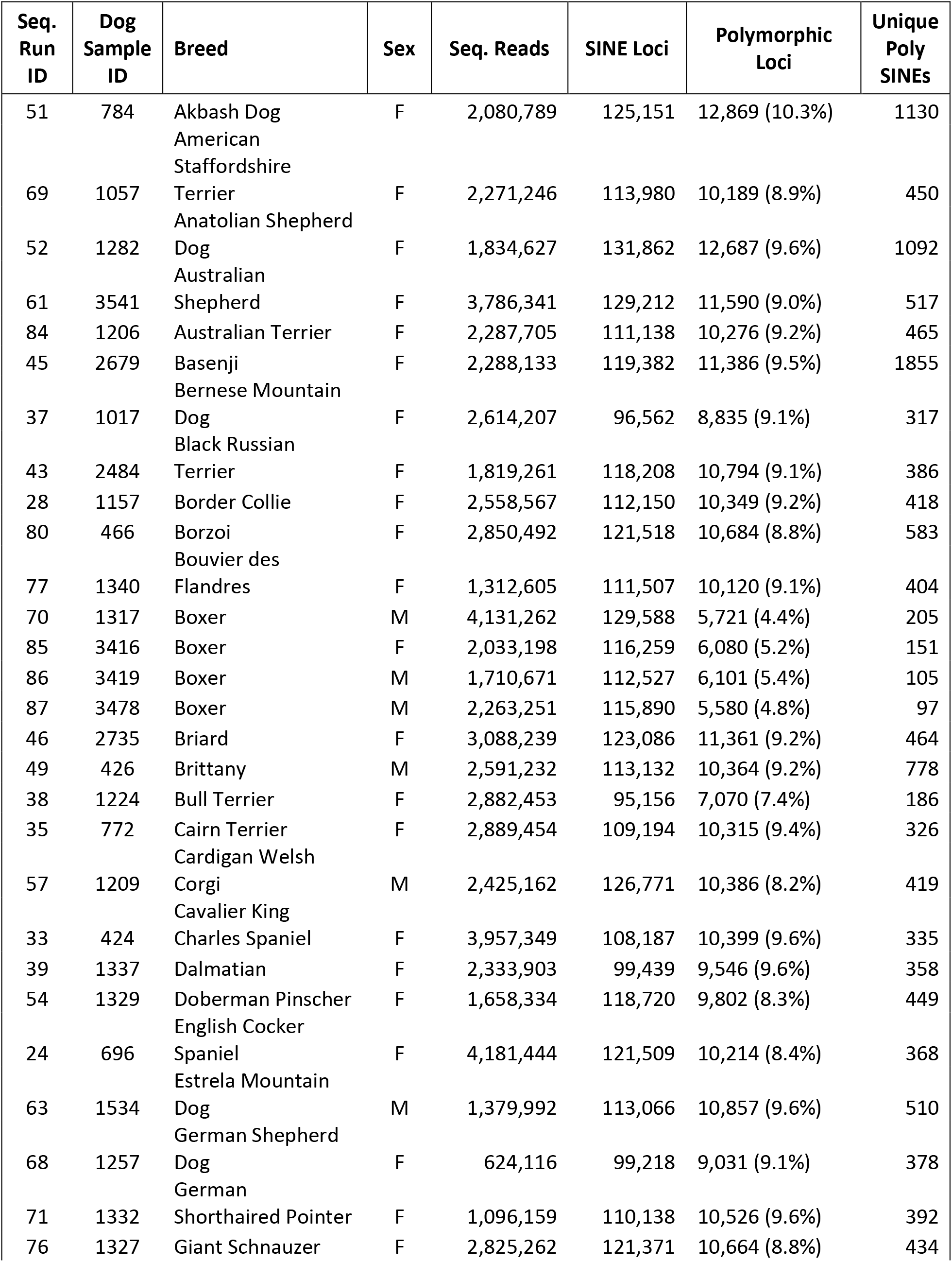
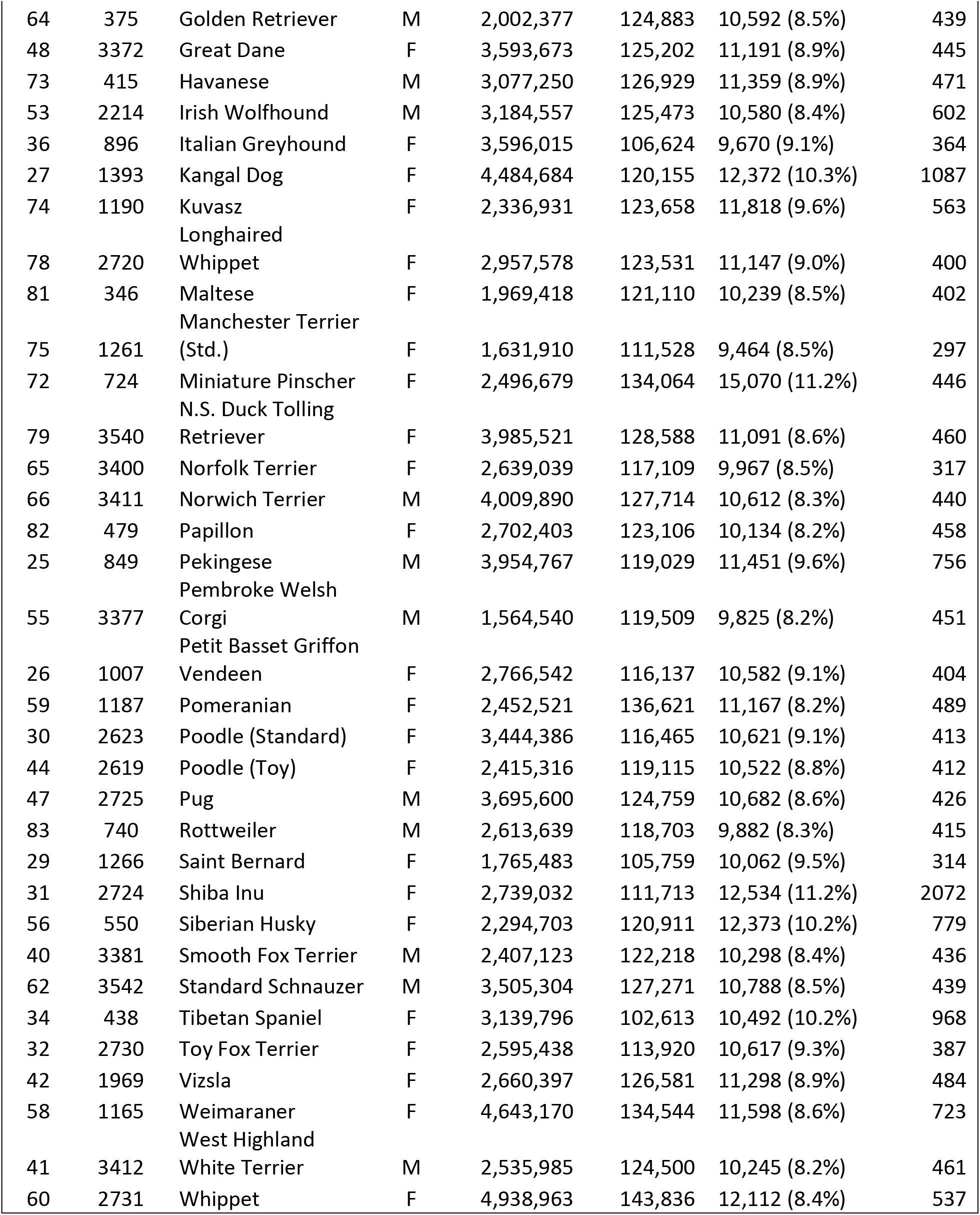
Dog libraries sequenced.

**Figure 2.**
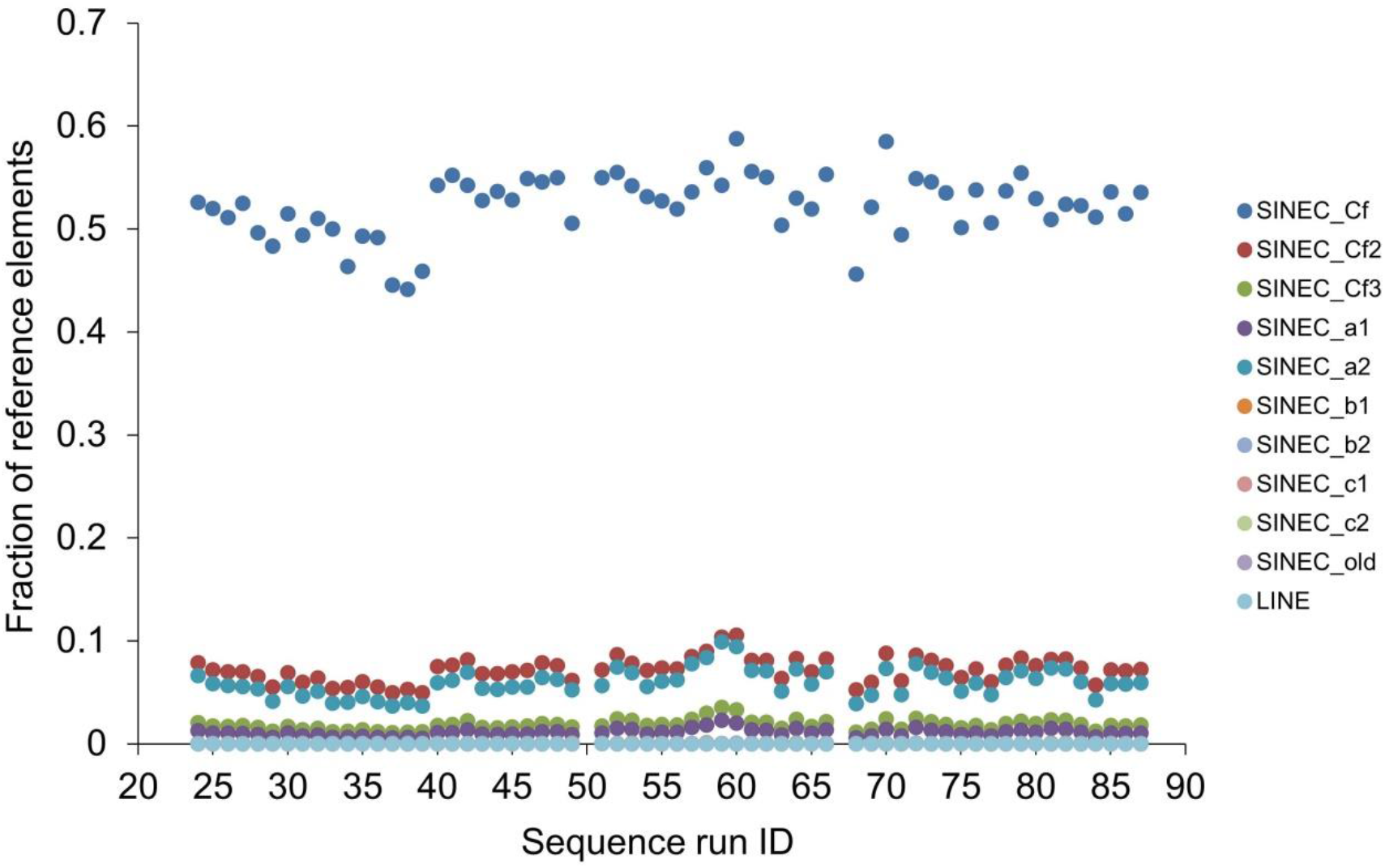
Libraries detect half of reference genome SINEC_Cf copies but few SINEs of other types.

Our libraries are specifically enriched for sequences flanking the SINEC_Cf sub-type of SINE. The libraries detect an average of just 7% of all reference SINEC_Cf2 copies and just 6% of all SINEC_a2 copies. Few SINEs of other sub-types and essentially no LINE copies are detected (Figure 2).

We define as polymorphic all SINE loci detected in one or more of our libraries but absent from the reference genome. Using this criterion, we identified 81,747 polymorphic SINEs in total (Table S2). An average of 13.3 reads support each polymorphic SINE but 11,193 SINE loci are called from just a single sequence read. Individual libraries detected between 5,580 and 15,070 polymorphic SINEs (mean=10,423) and these loci represent 4.4%-11.2% (mean=8.8%) of all SINE loci detected in a given library. The boxer libraries are low outliers (Table 1), probably due to our constraint that polymorphic SINEs be absent in the CanFam3 reference genome, which is from a boxer. Each library uniquely contributes an average of 518 SINE loci detected only in that library. The shiba inu and basenji contribute the largest numbers of unique SINE loci: 2072 and 1855, respectively. The akbash dog, Anatolian shepherd dog and Kangal dog each contribute more than 1000 unique SINE insertion loci. The four boxers and the closely related breed, bull terrier, contribute the smallest numbers of loci but in aggregate they add 744 unique SINE loci (Table 1).

To validate the polymorphic SINEs, we randomly selected 20 of them and PCR amplified the locus. We chose randomly selected DNA samples from among the dogs in which the library detected or did not detect the SINE insertion. We then genotyped the SINE insertions by assaying amplicon length on agarose gels. Three loci did not amplify, in four the SINE insertion was not confirmed and in the remaining 13 the insertion was supported. Our libraries detected three of the polymorphic SINEC_Cfs previously reported as causal for or associated with a trait selected by dog breeders (Table 2). These SINE polymorphisms have previously been detected in multiple breeds and the insertion allele has risen to high frequency in some breeds. We did not, however, detect any of the previously reported disease-causing SINE insertions (Table 2).

**Table 2.** Published polymorphic SINEs and LINEs causing or associated with traits in dog. ND=Not Determined because the published report does not precisely specify a genomic locus.

| FIRST AUTHOR | TRAIT | BREEDS | GENE | TYPE | WHERE | IN LIBS |
| --- | --- | --- | --- | --- | --- | --- |
| Mauldin | Ichthyosis | American Bulldog | NIPAL4 | SINE | Intergenic | ND |
| vonHoldt | Hypersociality | Many Breeds | Cfa6 locus | SINE | Several loci | ND |
| Goldstein | Retinal degeneration | Norwegian Elkhound | STK38L | SINEC_Cf | Coding Exon | No |
| Pele | Centronuc. Myopathy | Labrador Retriever | PTPLA | SINEC_Cf | Coding Exon | No |
| Weidmer | POANV | Alaskan Husky | RAB3GAP1 | SINEC_Cf | Coding Exon | No |
| Karlsson | Spotting | Boxer and others | MITF | SINEC_Cf | Intergenic | No |
| Sutter | Assoc. w/size | Small Breeds | IGF1 | SINEC_Cf | Intron | Yes; 38 |
| Dreger | Black and tan coat | Many Breeds | ASIP | SINEC_Cf | Intron, Invert. Rep. | Yes; 35 |
| Downs | Retinal atrophy | Tib. Spaniel & Terrier | FAM161A | SINEC_Cf | Intron, Near Exon | No |
| Lin | Narcolepsy | Doberman Pinscher | HCTR2 | SINEC_Cf | Intron, Near Exon | No |
| Clark | Merle coat | Herders | SILV | SINEC_Cf | Intron, Near Exon | Yes; 1 |
| Zeng | Bandera's ataxia | Coton de Tulear | GRM1 | LINE-1 | Coding Exon | No |
| Marchant | Brachyceph. | Bracycephalics | SMOC2 | LINE-1 | Intron | No |
| Smith | Muscular dystrophy | Pem. Welsh Corgi | DMD | LINE-1 | Intron | No |
| Brooks | hemophilia B | German Wire. Pointer | F9 | LINE-1 | Intron | No |

Using sizing of PCR amplicons we validated and genotyped 184 other SINE polymorphisms, including many in locations that have potential to alter gene expression or splicing (Table S3). We validated four polymorphic SINE insertions in coding exons. In *GSTA8P* we find just three out of 212 chromosomes have the insertion allele for a SINE in the 5^th^ to last codon. This mutation may not alter gene expression in any meaningful way, as the syntenic gene in human is listed as a pseudogene and is not expressed. We found a polymorphic SINE in *FANCD2’s* 2^nd^ to last coding exon in 12 chromosomes out of 63 dogs genotyped. Nine different breeds have the insertion allele and four genotyped dogs (Afghan hound, Chinese crested, Labrador retriever, and Italian greyhound) are homozygous for the SINE insertion. Splicing defects and other mutations in this gene cause Fanconi anemia in humans (Timmers *et al.* 2001). The *OXSM* gene has a SINE insertion in the stop codon (Figure 3) that is present in six different breeds. The insertion is homozygous in two Akitas and a miniature pinscher. In *SOX6* we found a coding exon SINE insertion only in four heterozygous dogs (Alaskan malamute, cavalier King Charles spaniel, Neapolitan mastiff, and saluki) out of the 63 genotyped animals from 41 breeds. The SINE is inserted in the downstream end of the 2^nd^ coding exon.

**Figure 3.**
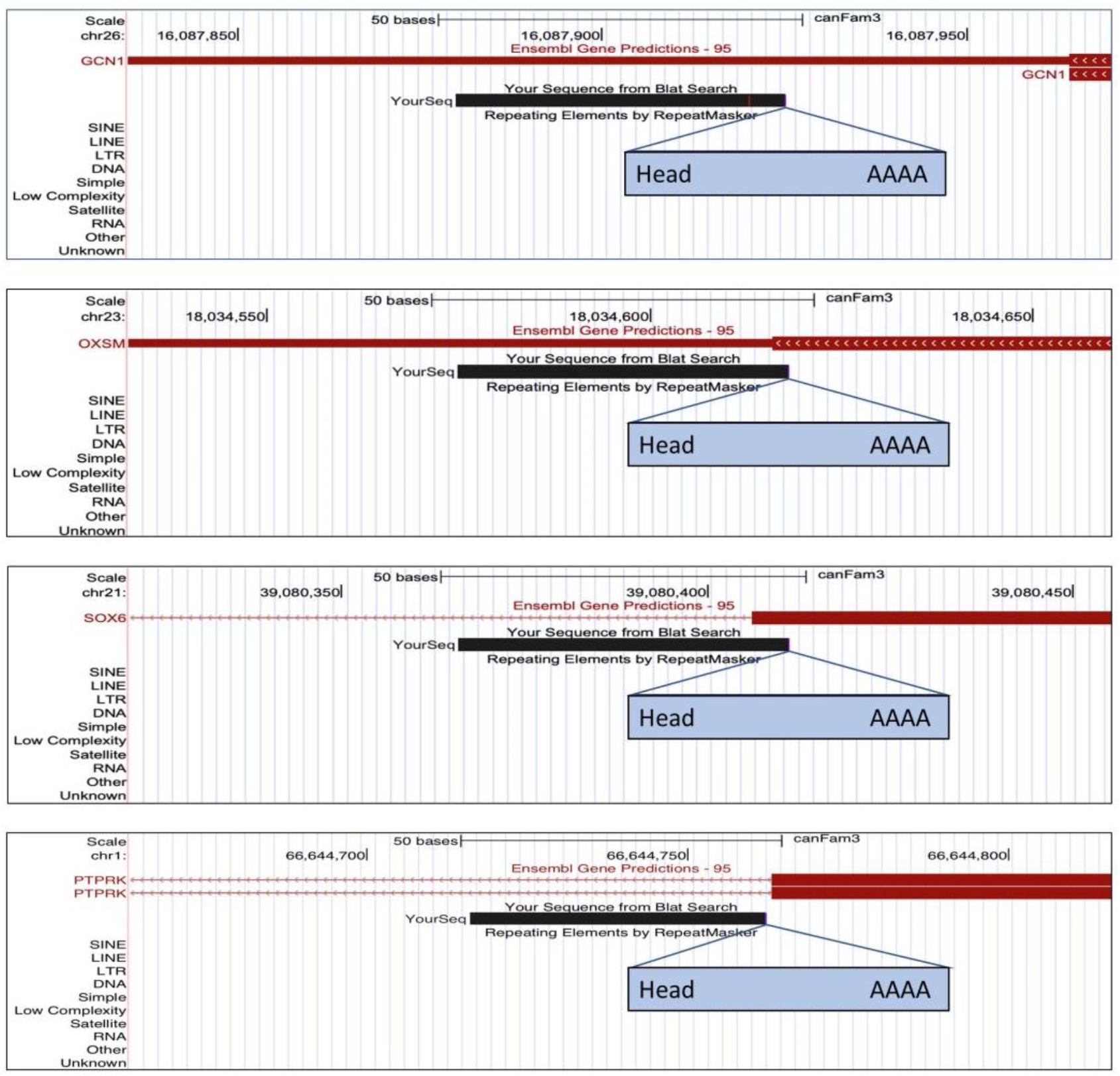
Examples of validated polymorphic SINE insertions in genes. The black bar indicates the “flank” portion of the Illumina sequence read that emerges from the head end of the SINE. There is a SINE insertion in the 3’UTR of *GCN1*. The polymorphic SINE in *OXSM* is located within the stop codon. *SOX6* has a SINE insertion located in the third coding exon. The SINE insertion in *PTPRK* is one bp downstream of coding exon 13.

We also validated SINE insertions in other genic contexts. We detected polymorphic SINEs in the 3’UTRs of *GCN1*, *KITLG*, *CAV3*, *DMD*, and ten other genes (Table S3). We validated SINEs in introns both upstream and downstream of nearby coding exons, including, for example, a SINE in sense orientation to *HMMR* 16 bp upstream of a coding exon. In *PLXNA2* we found an antisense SINE 23 bp upstream of the third coding exon. We validated 57 other intronic polymorphic SINEs that are not particularly close to coding exons, 27 in sense orientation and 30 in antisense orientation to the gene. We also validated 76 polymorphic SINEs in intergenic sequence (Table S3).

We initially investigated one SINE as a “near exon” type because it inserted just 44 bp upstream of a *WISP3* coding exon. But closer inspection revealed that a spliced expressed sequence tag (EST) from the downstream gene on the opposite strand, *TUBE1*, proceeds past *TUBE1’s* 3’UTR. The EST continues into *WISP3* and terminates precisely at the insertion site of the polymorphic SINE. It is intriguing to speculate that the SINE contributed to the EST’s termination although we don’t know the SINE genotype for the dog from which the EST was collected.

To put these SINE polymorphisms in a broader context we examined the patterns of SINE insertion into genes and intergenic sequence in the dog genome. The density of polymorphic SINEs is lowest in protein-coding exons. Density is comparable in intergenic and intronic sequences except that in introns the antisense orientation is favored over the sense orientation (Figure 4). In both intergenic and intronic sequences the autosomes have a higher density than chromosome X. Polymorphic SINE density is higher in 3’UTRs than 5’UTRs and there is no evidence for a strand bias. The boxer reference genome SINEC_Cf density patterns largely match the pattern in polymorphic SINEs except that chromosome X is not a low outlier for the reference genome SINEC_Cf density.

**Figure 4.**
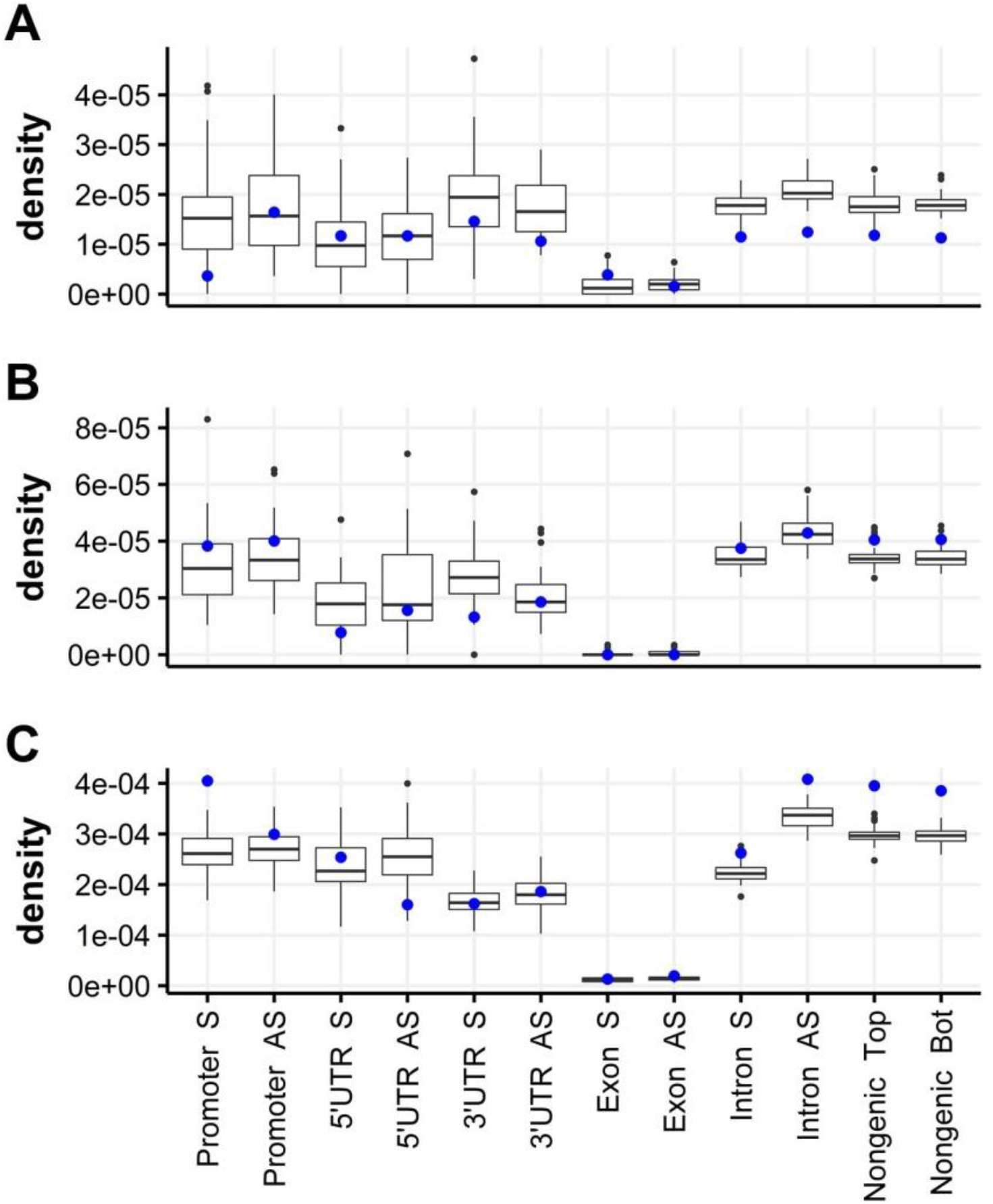
SINE and LINE density varies by strand and part of gene. Each boxplot shows the SINE or LINE density (the number of copies per ungapped bp) across the dog’s 39 chromosomes, with chr X specially indicated in blue. Gene regions followed by “S” show the density of sense strand SINEs or LINEs while “AS” shows the antisense density. (A) Polymorphic SINEs found with our libraries, (B) Reference genome SINEC_Cfs, and (C) Reference genome LINEs.

Other types of reference genome SINEs mostly match the pattern for SINEC_Cf (Figure S1). In most SINE types the intergenic density matches the sense-strand density within introns, however MIRs show no evidence for a strand bias in SINE density within introns. For all types of SINEs, presence in the coding exon is very rare. For LINEs (Figure 4C) there is an even greater strand bias in introns than that observed for SINEs, again with a higher density for antisense over sense-strand LINEs. LINEs are at a lower density in 3’UTRs compared to any other gene region except coding exons.

Polymorphic SINEs in certain gene regions are likely to be under negative selection due to their effects on gene expression and splicing patterns. We therefore hypothesized that polymorphic SINEs in certain genic contexts are likely to be found at low insertion frequencies. While an imperfect measure, we estimated SINE insertion allele frequency as the number of libraries in which we detected a given SINE. We also counted the number of polymorphic SINEs detected by just one library. For several different gene regions we find that about 40% of polymorphic SINEs are detected by just a single library (Table 3). For coding exons a larger share of all SINEs (61%) were detected just once, consistent with the idea that these SINEs have a lower than average insertion allele frequency. Furthermore, the coding exon polymorphic SINEs are found in just four libraries on average, below the averages for other genic regions. Although not a significant difference after correcting for multiple hypothesis testing, we find a trend to a lower number of detection libraries for polymorphic SINEs close to exons as compared to intronic SINEs as a whole.

**Table 3.**
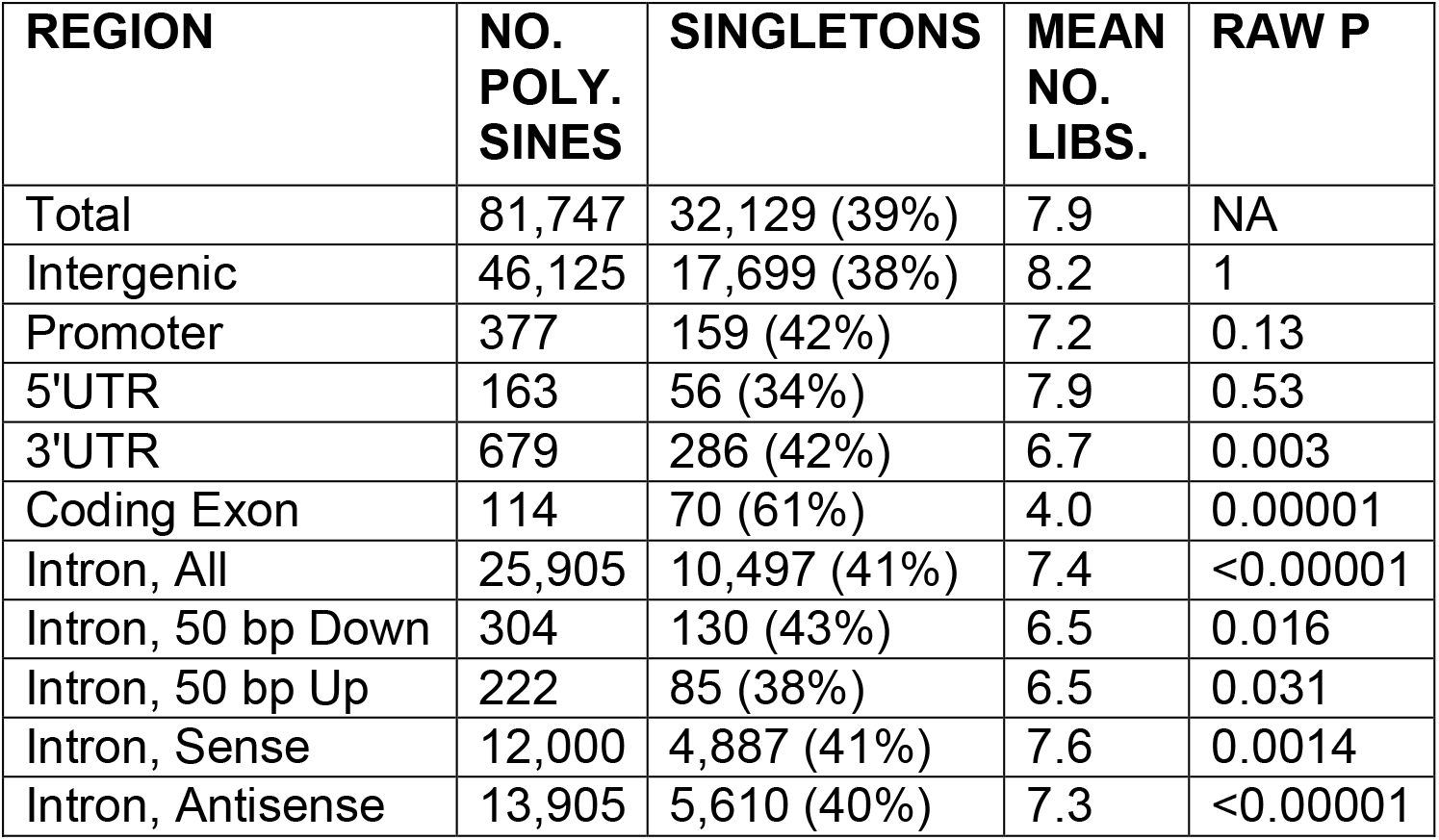
SINE insertion frequencies.

We inspected all 83 putative polymorphic SINEs intersecting with coding exons (Table S4). Based on synteny to the human genome and an absence of introns we scored 23 SINEs (28%) as likely to be in pseudogenes. The remaining 60 SINEs are inserted into coding exons of genes with synteny to the human reference. For many of these genes mutation in the human or mouse leads to disease or other traits. For example, the American Staffordshire terrier, great Dane, and Vizsla breed libraries discover a SINE insertion in *POGLUT1*. When the ortholog is knocked out in the mouse there is embryonic lethality and large disruptions in development of the nervous system (Ramkumar *et al.* 2015). As is the case for many of our exonic insertions, the entire SINE-flanking sequence is located within the exon which leaves open the possibility that the SINE is instead inserted into a retrogene of *POGLUT1* present in our library dogs but absent from the dog reference genome. An insertion detected in three of our libraries is in exon 7 of *TEX15*. Human males homozygous for mutation in this gene are infertile (Okutman *et al.* 2015). None of our library dogs were phenotyped for fertility and from library sequencing alone we can’t assess whether any are homozygous for the SINE insertion. *USP2* mutation reduces male fertility in mice (Bedard *et al.* 2011); we found a SINE insertion in exon 7 of *USP2* in our German shorthaired pointer library.

In 28 libraries we detected an insertion into the *SFPQ* gene. *SFPQ* helps ensure RNA pol II progresses through long introns of genes expressed in the human brain (Takeuchi *et al.* 2018). An insertion into the *IGHM* gene is detected in two of our libraries. This gene is part of the constant region of immunoglobulin heavy chains and a human family with a mutation in exon 1 segregates agammaglobulinemia (Silva *et al.* 2017). As a final example, in 11 of our libraries we detect a SINE insertion in the first coding exon of *GJE1*, a connexin component of gap junction channels that plays a role in hearing (Yang *et al.* 2005).

We next investigated SINE and LINE insertions in introns. The strand bias for intronic reference SINEC_Cf and LINE copies is a general feature of introns along their length (Figure 5) and the density of both types of retrotransposons declines near exons both upstream and downstream. Reference genome SINEs and LINEs are likely to be fixed and therefore, on average, are older than polymorphic SINEs that have not yet gone to fixation (or not yet been removed from the population). We next asked whether the density of polymorphic SINEs also drops near exons. Downstream of exons we see no drop in density, even close to exons (Figure 6). However, upstream of exons there is a drop in polymorphic SINE density for both sense and antisense SINEs. The density decrease is evident only for the first few tens of bp for both orientations.

**Figure 5.**
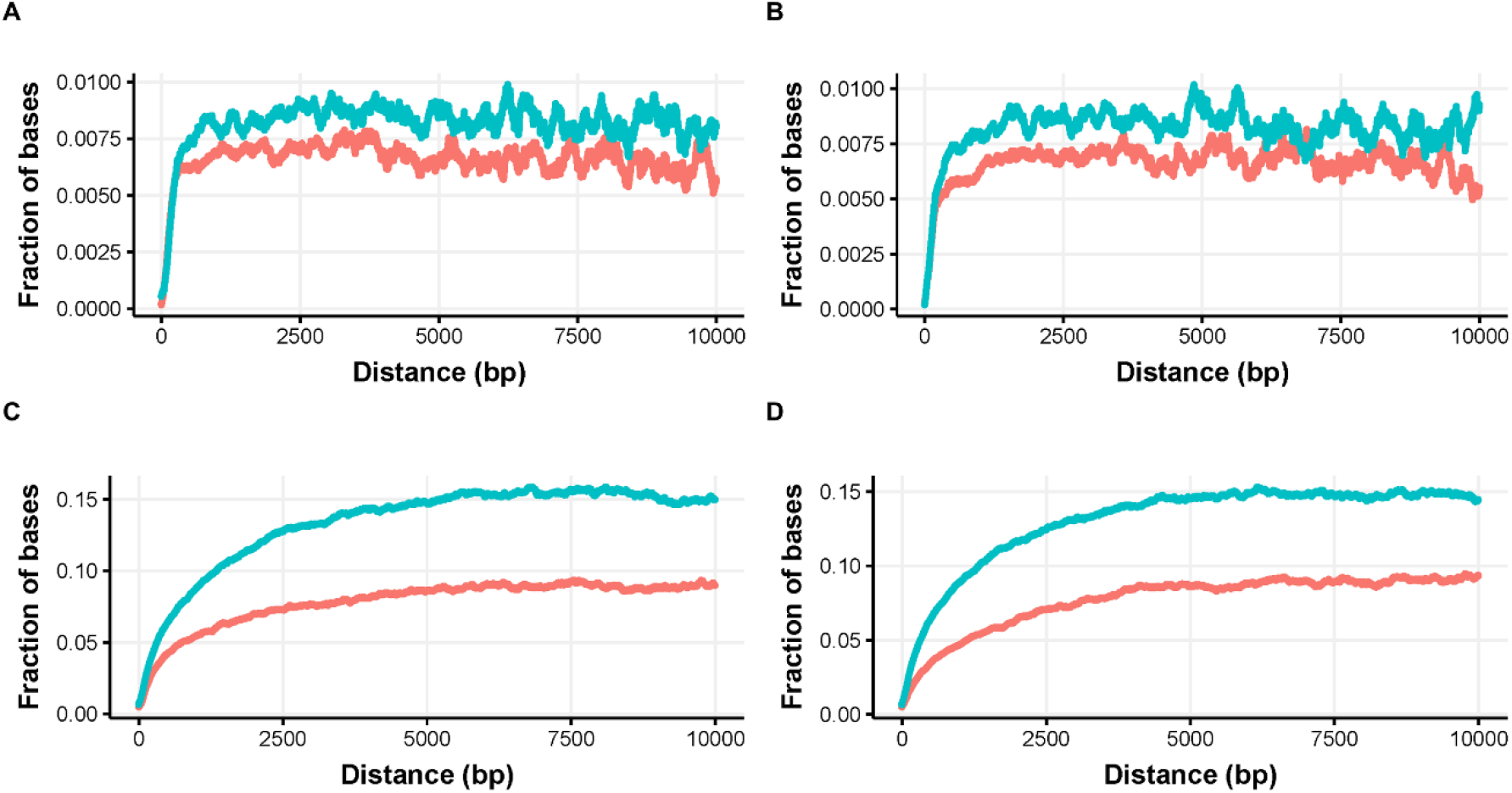
Reference SINEC_Cfs and LINEs in introns have a strand bias and their density decreases near exons. The y axis displays the fraction of introns in which the bp is retrotransposon. The red curve is sense strand and the blue curve is antisense strand. (A) SINEC_Cf in introns upstream from an exon, (B) SINEC_Cf in introns downstream from an exon, (C) LINEs in introns upstream from an exon and, (D) LINEs in introns downstream from an exon.

**Figure 6.**
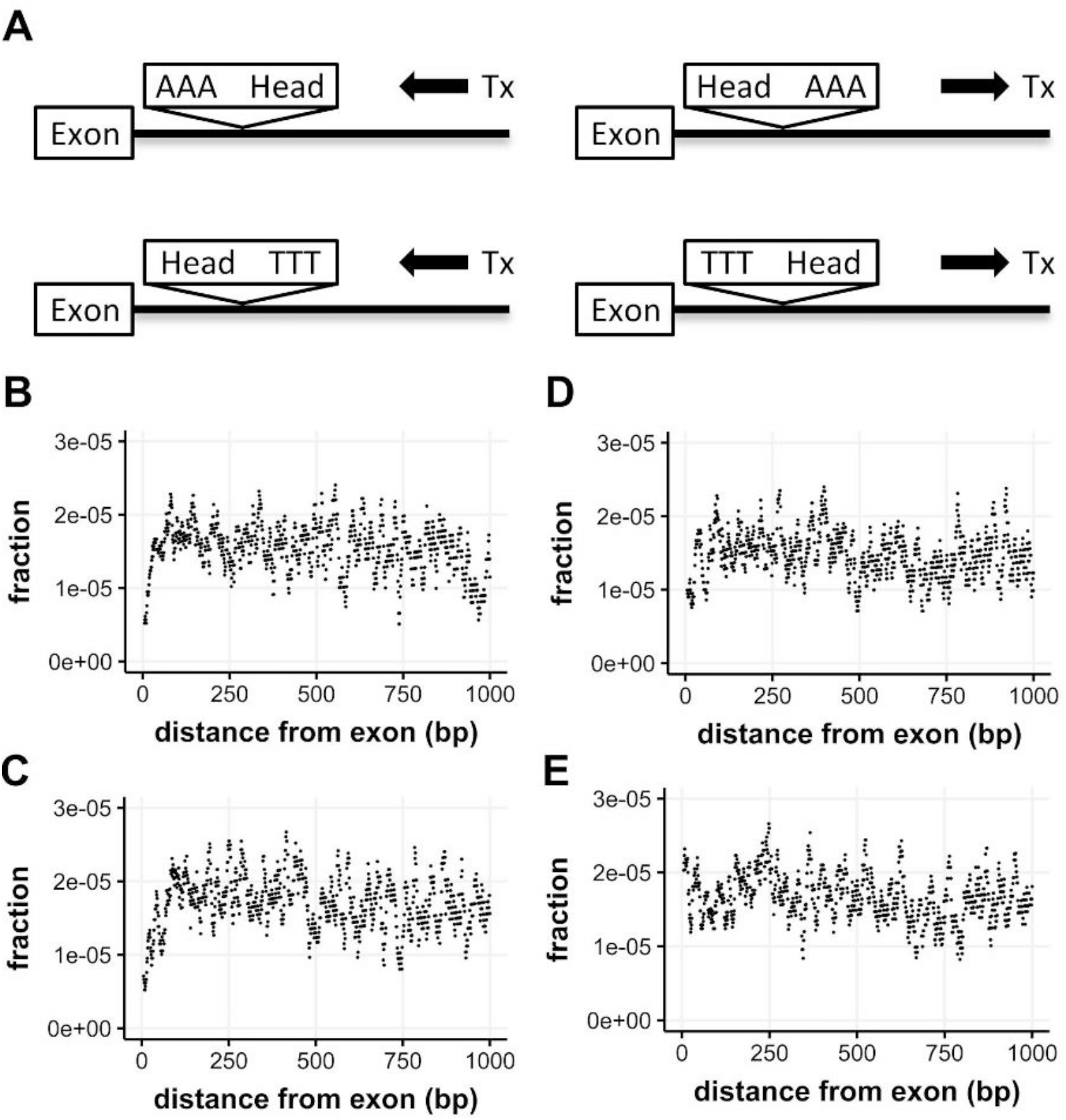
Polymorphic SINEs in introns are rarely found upstream of nearby exons. (A) The four possible orientations of SINE insertions were tracked. Tx = direction of gene transcription. (B-E) The y axis plots the fraction of all dog genome intronic bases x distance from an exon. The value plotted on the x axis is the center of a moving average +/− 5 bp. (B) Distance up into introns towards the 5’ end of the gene; the polymorphic SINE is in the sense orientation, as shown in the top left of panel A. (C) Distance up into introns; polymorphic SINEs are in antisense orientation. (D) Distance down into introns; polymorphic SINEs are in sense orientation. (E) Distance down into introns; polymorphic SINEs are in antisense orientation.

We next asked whether the dog genome contains evidence of bias against inverted pairs of retrotransposons, as has been found in other mammals (Cook *et al.* 2011), where inverted pairs are rare compared to direct repeats. We therefore enumerated all cases in the dog reference genome introns in which reference SINEC_Cf pairs exist within 1000 bp of each other as nearest neighbors (Figure 7). Pairs within about 75 bp of each other have a marked bias where both types of inverted pairs, head-tail and tail-head, are rare compared to both types of direct repeats, head-head and tail-tail. This bias has the same strength and directionality for SINE pairs in intergenic sequence as well (Figure 7B). Furthermore, for both introns and intergenic sequences the bias is correlated with the strength of the pairwise sequence alignment score between the two SINEC_Cf copies. The dog genome contains very few cases of SINEC_Cfs with nearly identical sequence close together in inverted orientation.

**Figure 7.**
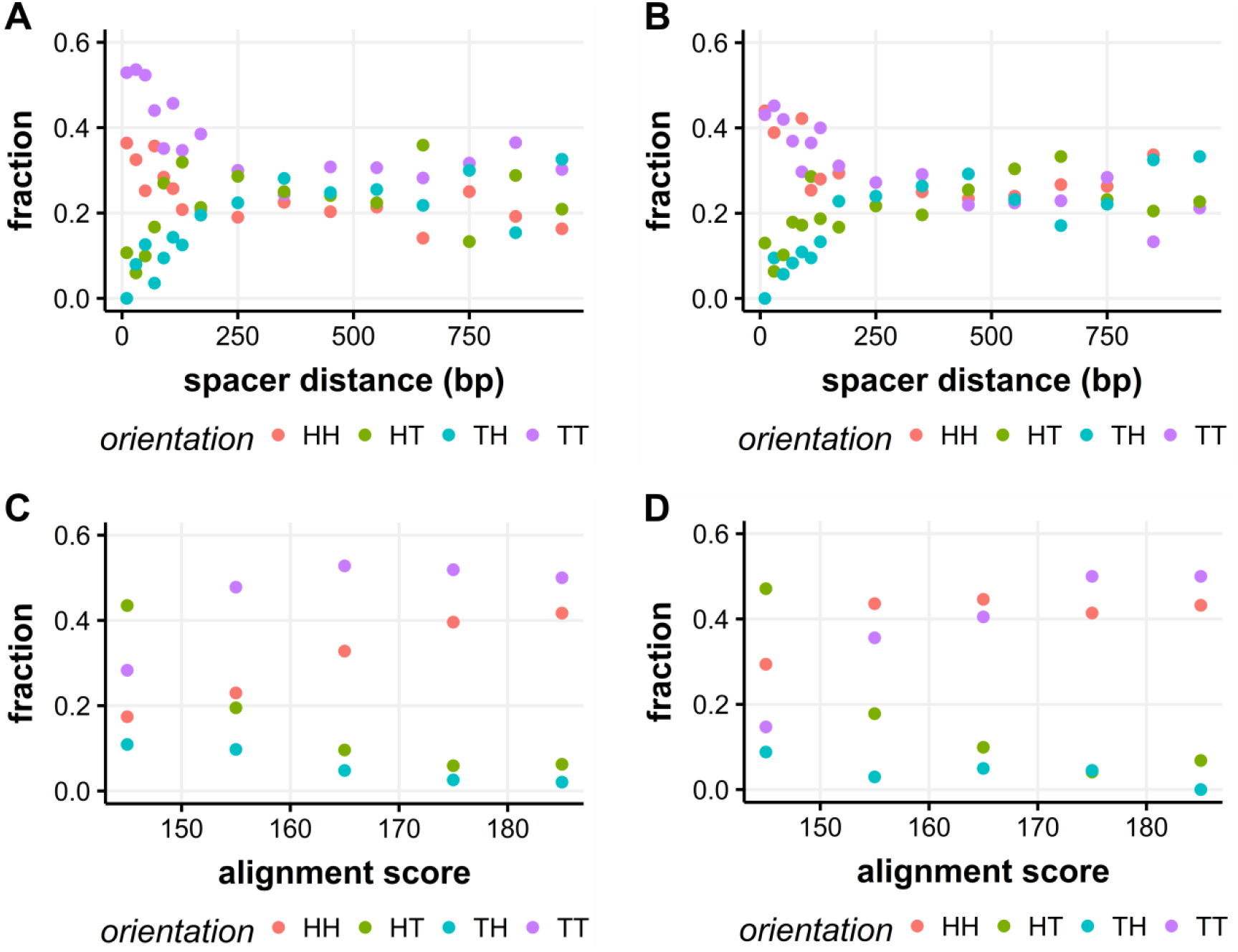
Inverted pairs of reference SINEC_Cf copies are rare when the SINEs are close and similar in sequence. (A) The four possible orientations of pairs in introns were tracked relative to the gene. HH = both SINEs are in sense orientation (RNA pol encounters the heads first); TT = both antisense; HT = RNA pol hits the head of the first SINE then the tail of the second; TH = RNA pol hits the tail of the first SINE then the head of the second. {HH,TT} are direct pairs while {HT, TH} are inverted pairs. (B) Intergenic pairs, H = top strand, T = bottom strand. All intronic (C) and intergenic (D) pairs with spacers up to 100 bp were included.

LINE pairs in the dog genome also show a bias against inverted pairs but the pattern is different from SINEC_Cfs (Figure 8). For LINE pairs the bias holds even when the LINEs are separated by as much as ~500 bp apart, perhaps as much as 10 times greater distance than for SINEC_Cf. As with SINEs, the LINE bias is observed in both introns and intergenic sequences. Unlike for SINEs, the LINEs show less evidence for pairwise alignment score positively correlating with bias against inverted pair orientations. In introns, LINE pairs with highly similar sequence are much more likely to both be antisense (tail-tail) than any of the other three possible orientations (Figure 8C).

**Figure 8.**
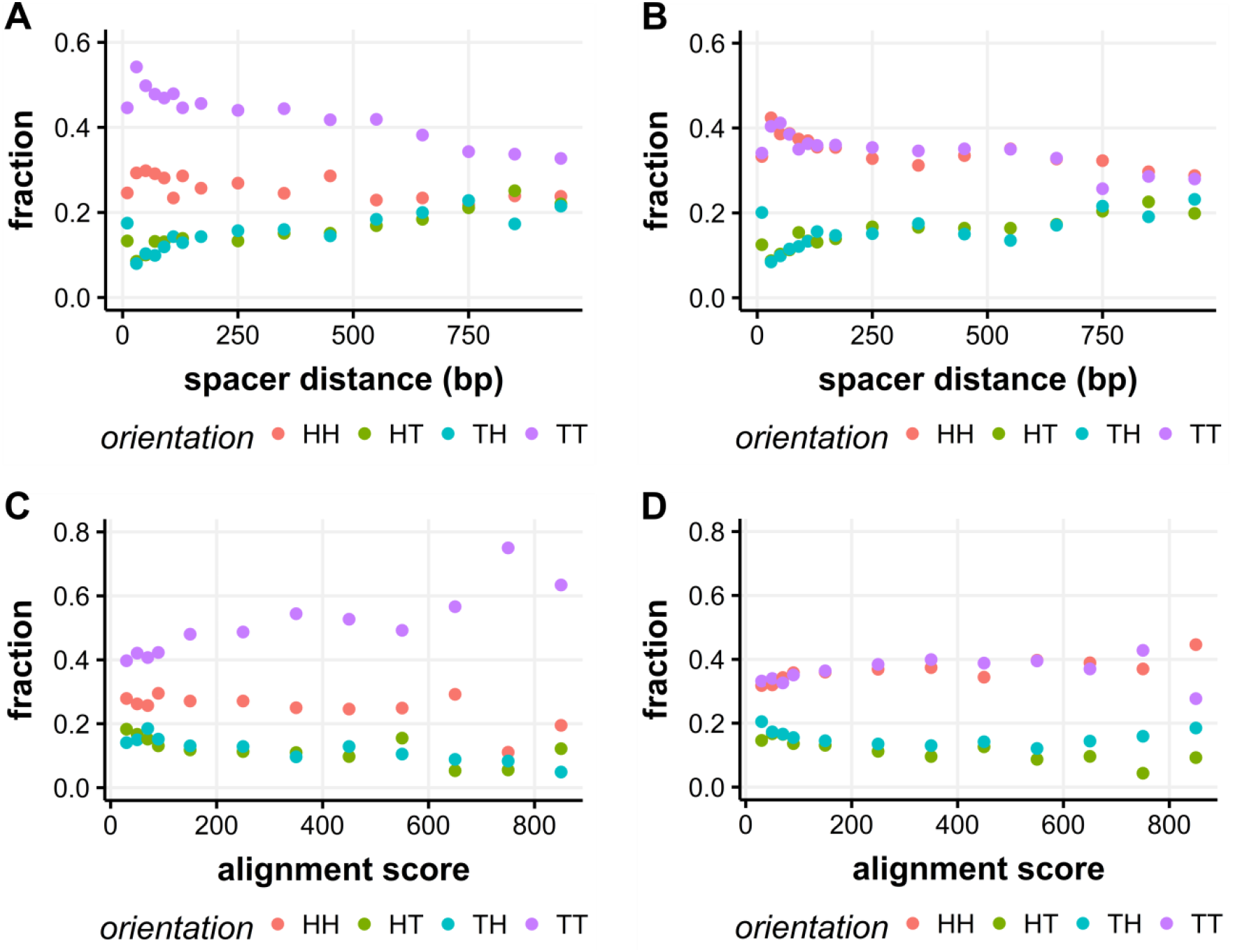
Inverted pairs of reference LINE copies are rare when the LINEs are close and similar in sequence. (A) The four possible orientations of pairs in introns were tracked relative to the gene. HH = both LINEs are in sense orientation (RNA pol encounters the heads first); TT = both antisense; HT = RNA pol hits the head of the first LINE then the tail of the second; TH = RNA pol hits the tail of the first LINE then the head of the second. {HH,TT} are direct pairs while {HT, TH} are inverted pairs. (B) Intergenic pairs, H = top strand, T = bottom strand. All intronic (C) and intergenic (D) pairs with spacers up to 500 bp were included.

With evidence that reference SINEC_Cf pairs are rarely inverted when close together, we hypothesized that our polymorphic SINEs would show the same bias against inverted pairs when their closest SINE neighbor was SINEC_Cf. Because our sequence reads emanating from and flanking the head end of each SINE are only ~65 bp, the presence of repeated sequence in them strongly impacts confident alignment to the genome. We therefore filtered out sequence reads containing too much repeated sequence (see methods). Consequently, tail-head inverted pairs that are very close to one another are undercounted since a repeat (the reference SINE) would be found in the head-flanking sequence of our polymorphic SINE and this would cause that polymorphic SINE’s sequence reads to be excluded from our dataset. We thus assayed polymorphic SINE pair bias using only the head-tail type of inverted pairs plus both types of direct repeats. At very close distances, roughly 0-50 bp apart, there is a lack of head-tail inverted pairs compared to direct repeats (Figure 9). This is evident in both introns and intergenic sequence and the bias is present both when our polymorphic SINE is close to a reference SINEC_Cf or close to a SINEC_Cf2. However, no bias is found for polymorphic SINEs close to either LINEs or MIR-type SINEs (Figure 9E-H).

**Figure 9.**
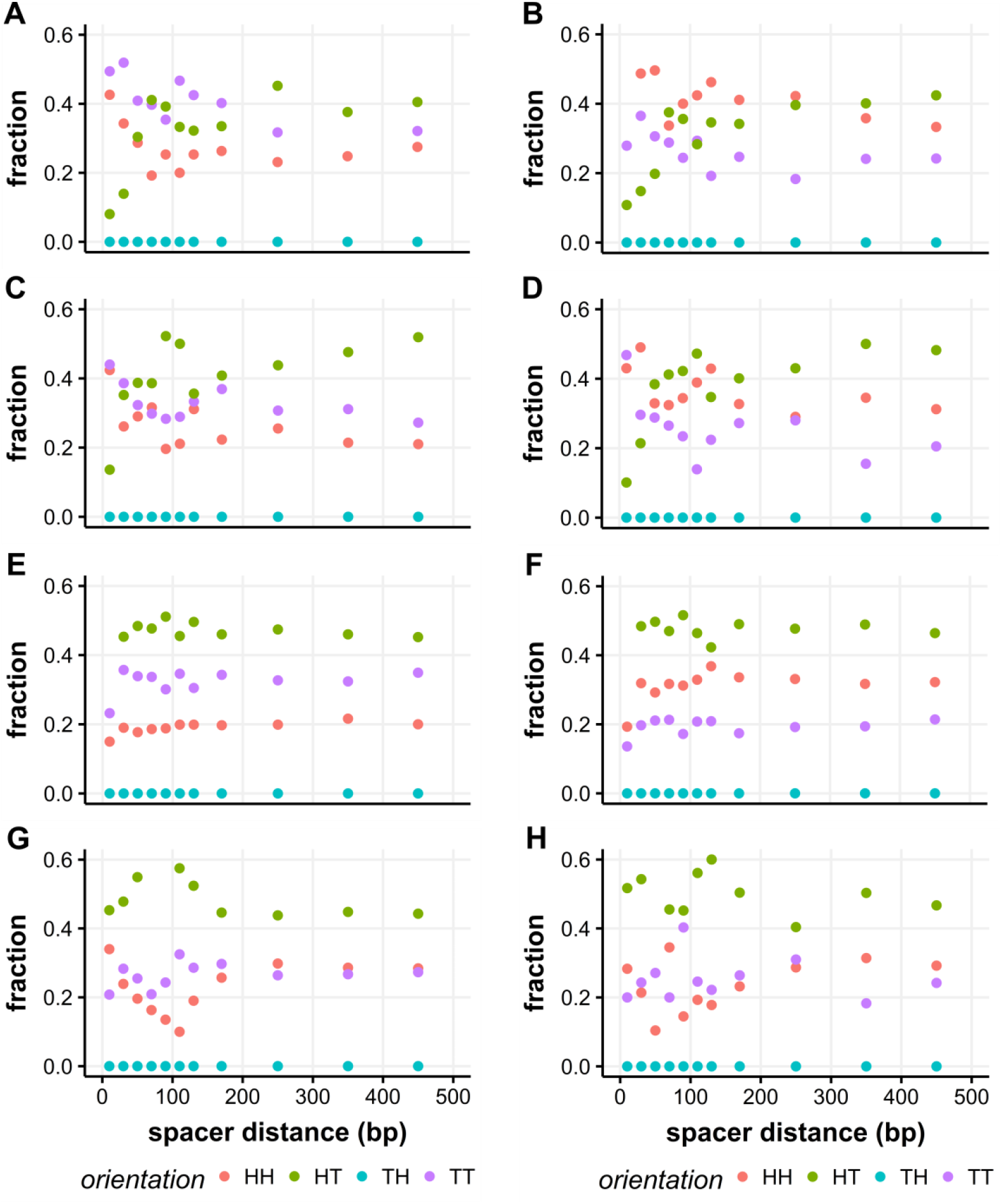
Polymorphic SINEs near SINEC_Cf and SINEC_Cf2 are rarely in inverted orientation. Because head-proximal flanks containing repeat sequences were filtered out of our dataset during SINE discovery (see methods) we had to exclude from the analysis all pairs in which the polymorphic SINE’s head faces the reference SINE. Thus the “TH” orientation can’t be found. The three remaining orientations of pairs were tracked in relation to the gene’s direction of transcription (left column) or top strand (right column). HH = both in pair are in sense orientation; TT = both antisense; HT = heads face out (inverted orientation). Polymorphic SINEs are paired with: (A) reference SINEC_Cf in introns and (B) intergenic sequence, (C) SINEC_Cf2 in introns and (D) intergenic, (E) LINEs in introns and (F) intergenic sequence, and (G) MIRs in introns and (H) intergenic sequence.

## Discussion

The dog population carries in its many genome copies a tremendous resource: a natural experiment in mutagenesis via retrotransposition. It is clear that the dog genome carries many thousands of young polymorphic SINE insertions (Kirkness *et al.* 2003; Lindblad-Toh *et al.* 2005; Wang and Kirkness 2005). SINE insertions in other mammals disrupt gene expression and splicing in many ways: untranslated region insertion, coding exon insertion, intronic insertions near or far from exons, and via interaction with a second SINE in inverted orientation (Cook *et al.* 2011; Taskesen *et al.* 2012; Anwar *et al.* 2017).

In the dog, genome-scale analysis of SINE molecular genetics lags behind other mammals like human and mouse. However, by mapping both selected and disease traits in the dog, researchers have over the past 20 years built a long list of phenotypes attributable to SINEC_Cf insertion. Polymorphic insertions of both SINEC_Cf and LINE1 cause phenotypes in dog breeds. We are not aware of LINE1 insertions that are positively selected for, but SINEC_Cf insertions cause phenotypes that range from serious diseases like narcolepsy (Lin *et al.* 1999) and retinal atrophy (Goldstein *et al.* 2010; Downs and Mellersh 2014) to the positively selected coat patterns merle (Clark *et al.* 2006; Murphy *et al.* 2018) and black-and-tan (Dreger and Schmutz 2011). It would be interesting to scan dog genomes for polymorphic LINE1 insertions like we’ve done here with SINEC_Cf. We speculate that polymorphic LINE1 insertions have also contributed to the great trait diversity in dogs.

Three traits, namely narcolepsy in Doberman pinschers, retinal atrophy in Tibetan spaniels and terriers, and merle coat in Australian shepherds and other breeds are all caused by SINEC_Cf insertion within an intron near to, and upstream of, a coding exon. With merle the SINE is immediately within the exon/intron junction but with narcolepsy the SINE is 35 bp upstream. In all three cases the SINE is in antisense orientation. While a sample of three mutations is very small, a pattern is suggested. By looking at every Ensembl gene model in the dog canFam3.1 reference genome we find a low density of polymorphic SINEC_Cfs upstream, but not downstream, of coding exons. This is consistent with the idea that such a location is likely to alter gene function, causing these insertions to be removed from the population via purifying selection. Although we don’t favor the hypothesis, our data don’t exclude the possibility that SINE insertion upstream of coding exons occurs at a lower rate than other genic or intergenic locations. Our data fit with the mapped trait-causing SINEs well. Because we found low SINE densities not just for antisense but also for sense-oriented SINEs we predict that sense-oriented SINEC_Cf insertions upstream of coding exons are also capable of causing disease or other phenotypes. Any polymorphic SINE immediately upstream of a coding exon thus warrants extra scrutiny. Because we found the density of our polymorphic SINEs is normal immediately downstream of coding exons perhaps we should expect few or no trait-causing SINE insertions to be reported in the future in those positions. That said, we do find a “hazardous zone” (Zhang *et al.* 2011) of very low density for reference (mostly fixed) SINEC_Cf copies surrounding coding exons both upstream and downstream.

It may be that SINEC_Cf insertion near an exon is generally deleterious but selection against upstream events is stronger and provides a more obvious signal that is detectable with our polymorphic SINE dataset. If upstream insertions are more strongly deleterious than downstream insertions it explains the fact that near-exon dog disease-causing SINEC_Cf insertions are reported for upstream positions. Comparison of gene expression datasets with patterns of polymorphic SINEs could distinguish the potential roles of upstream vs. downstream near-exon SINE insertions.

SINEs can function within genomes in many different ways. The genomes of human and mouse have been shown to carry biased ratios of SINE pairs. While a null model would predict head-head, tail-tail, head-tail and tail-head orientations of pairs should all occur at equal frequency, what is seen instead is a lack of the inverted pair orientations: head-tail and tail-head. In these orientations the SINE transcripts are opposite strands and capable of hybridizing with each other. The loss of inverted pairs is particularly evident with a short spacer distance separating the two SINEs, although altered ratios of direct:inverted repeats have been reported for huge spacer distances (Cook *et al.* 2013). We observe altered ratios of direct:inverted repeats for pairs of reference SINEC_Cfs and LINEs. The majority of reference SINEs and LINEs are presumably fixed so the pairs are a feature of dog genomes in general. We detect a fairly short-range ratio bias for SINEC_Cf pairs; the spacer distance may be no more than approximately 100 bp. While we don’t find evidence for biased ratios at greater distance this may be due to our approach. Unlike analyses in the human genome, here we only examined the pairs of adjacent nearest neighbor SINEs rather than all combinations of pairs within some interval. We find indirect support for the idea that transcripts of the two SINEs within pairs are hybridizing: the loss of inverted SINE pairs is correlated with the pairwise sequence alignment score of the two SINEs. Thus, few SINEC_Cf inverted pairs in the reference genome are close together and of those that are, very few have high sequence identity.

Most SINEs that compose pairs in the reference genome are likely to be fixed. Therefore, it seems likely that at least some of these rare, inverted, closely-opposed pairs may have functional roles to alter gene expression in adaptive ways. Searching for these pairs in wild canids like the gray wolf and coyote could help put bounds on their ages. We found that short-spacer inverted SINE pairs are underrepresented in intergenic sequence, just as they are in introns. This matches findings in other mammals (Lev-Maor *et al.* 2008; Cook *et al.* 2011).

We find that both LINEs and SINEC_Cfs have a higher density in dog introns when in the antisense rather than sense orientation. This bias is particularly marked for LINEs: sense-oriented intronic LINEs have a lower density than intergenic LINEs but antisense-oriented LINEs have a higher density than intergenic LINEs. Similarly, human *Alu* and mouse B SINEs have higher antisense than sense density in introns (Tsirigos and Rigoutsos 2009). As seen in other mammals, in dogs the density of LINEs in intergenic and intronic sequence is also higher on the X chromosome than autosomes (Bailey *et al.* 2000). New L1s in human may be targeting chromosome 1 rather than X (Sultana *et al.* 2019) and it would be interesting to know if the same is happening in dogs.

Here we report a large collection of polymorphic SINEC_Cf retrotransposon insertions in the dog genome. In our libraries on average 9% of all SINE loci are judged polymorphic because they are absent from the reference boxer. This value closely matches the 8% of polymorphic SINEC_Cfs reported for the boxer reference genome (Lindblad-Toh *et al.* 2005). This is a high proportion of all loci and suggests that many young SINE insertions are segregating across purebred dog populations. We found and validated many dozens of SINEs inserted into genes in ways that, in other genes at least, lead to disruption of gene expression or splicing patterns. We focused on SINEC_Cf because it is a young SINE and previous work has shown that many thousands of insertions are segregating in the dog genome (Kirkness *et al.* 2003; Wang and Kirkness 2005). Because our libraries can be pooled at 16-plex they represent a relatively inexpensive way to discover many thousands of SINEs. However, our strategy is limited in several ways. First, we require hybridization of a primer in the SINE sequence. This young dog SINE is easy to differentiate from older SINEs (Wang and Kirkness 2005) but there is no guarantee that primer hybridization would work so well in other species. Second, our library production includes several steps that require cycles of PCR so there is potential to favor some loci over others. Third, we collected and aligned short reads; our locus determination is based on alignment of ~60 bp SINE flanks to the genome. Mappability of these shorter reads, due to the presence of repeat sequences or other factors, limits our detection of some SINEs. For these reasons, despite the fact that we discovered tens of thousands of SINEs, our catalog of SINEs is not exhaustive. Future analysis of SINE insertions into SINEs or other repeat sequences may be a particularly fertile ground for discovery of new polymorphic SINEs. SINE insertion into a SINE or LINE poly-A tail is likely to be common in the dog genome but, because of our short flanking sequence reads, is not something our experimental design is well-suited to address. Cataloging SINE insertions within retrotransposons could help elucidate the role of SINE pairs in dog gene expression.

Just as *Alu* insertions can cause disease in humans, SINEC_Cf insertions sometimes cause disease in dogs. SINE insertion alleles in dogs have also, sometimes, been selected for by humans. The merle and black-and-tan SINE alleles have risen to high frequency in several breeds. Although most SINE insertions that alter dog genes are likely to be deleterious, we hypothesize that adaptive SINE insertions also exist and await discovery. Characterizing SINE insertions puts us one step closer to a comprehensive understanding of the dog genome.

## Acknowledgments

We thank David Kupiec, Florencia Ardon and Lalit Ponnala for technical assistance, and Peter Schweitzer at the Cornell University Life Sciences Core Lab for help troubleshooting barcoding issues with our custom Illumina HiSeq libraries. We also thank the dog owners and breeders who contributed samples from their animals. Research reported in this publication was supported by the National Human Genome Research Institute of the National Institutes of Health under award number R21HG006051. The content is solely the responsibility of the authors and does not necessarily represent the official views of the National Institutes of Health. This research was also supported by internal funds from Cornell University and La Sierra University.

## Literature Cited

Adelson, D. L., J. M. Raison, and R. C. Edgar, 2009 Characterization and distribution of retrotransposons and simple sequence repeats in the bovine genome. PNAS 106: 12855–12860.

Anwar, S. L., W. Wulaningsih, and U. Lehmann, 2017 Transposable Elements in Human Cancer: Causes and Consequences of Deregulation. International Journal of Molecular Sciences 18: 974.

Bailey, J. A., L. Carrel, A. Chakravarti, and E. E. Eichler, 2000 Molecular evidence for a relationship between LINE-1 elements and X chromosome inactivation: The Lyon repeat hypothesis. Proc Natl Acad Sci U S A 97: 6634–6639.

Bedard, N., Y. Yang, M. Gregory, D. G. Cyr, J. Suzuki et al., 2011 Mice Lacking the USP2 Deubiquitinating Enzyme Have Severe Male Subfertility Associated with Defects in Fertilization and Sperm Motility. Biology of Reproduction 85: 594–604.

Boyko, A. R., P. Quignon, L. Li, J. J. Schoenebeck, J. D. Degenhardt et al., 2010 A Simple Genetic Architecture Underlies Morphological Variation in Dogs. PLOS Biology 8: e1000451.

Chen, L.-L., and G. G. Carmichael, 2008 Gene regulation by SINES and inosines: biological consequences of A-to-I editing of Alu element inverted repeats. Cell Cycle 7: 3294–3301.

Chernova, T., F. M. Higginson, R. Davies, and A. G. Smith, 2008 B2 SINE retrotransposon causes polymorphic expression of mouse 5-aminolevulinic acid synthase 1 gene. Biochemical and Biophysical Research Communications 377: 515–520.

Clark, L. A., J. M. Wahl, C. A. Rees, and K. E. Murphy, 2006 Retrotransposon insertion in SILV is responsible for merle patterning of the domestic dog. PNAS 103:1376–1381.

Coltman, D. W., and J. M. Wright, 1994 Can SINEs: a family of tRNA-derived retroposons specific to the superfamily Canoidea. Nucl Acids Res 22: 2726–2730.

Cook, G. W., M. K. Konkel, J. D. Major, J. A. Walker, K. Han et al., 2011 Alu pair exclusions in the human genome. Mobile DNA 2: 10.

Cook, G. W., M. K. Konkel, J. A. Walker, M. G. Bourgeois, M. L. Fullerton et al., 2013 A Comparison of 100 Human Genes Using an Alu Element-Based Instability Model (G. Ast, Ed.). PLoS ONE 8: e65188.

Daniel, C., M. Behm, and M. Öhman, 2015 The role of Alu elements in the cis-regulation of RNA processing. Cell. Mol. Life Sci. 72: 4063–4076.

Das, M., L. L. Chu, M. Ghahremani, T. Abrams-Ogg, M. S. Roy et al., 1998 Characterization of an abundant short interspersed nuclear element (SINE) present in Canis familiaris. Mammalian Genome 9: 64–69.

Downs, L. M., and C. S. Mellersh, 2014 An Intronic SINE Insertion in FAM161A that Causes Exon-Skipping Is Associated with Progressive Retinal Atrophy in Tibetan Spaniels and Tibetan Terriers (C. Wade, Ed.). PLoS ONE 9: e93990.

Dreger, D. L., and S. M. Schmutz, 2011 A SINE Insertion Causes the Black-and-Tan and Saddle Tan Phenotypes in Domestic Dogs. Journal of Heredity 102: S11–S18.

Gagnier, L., V. P. Belancio, and D. L. Mager, 2019 Mouse germ line mutations due to retrotransposon insertions. Mobile DNA 10: 15.

Goldstein, O., A. V. Kukekova, G. D. Aguirre, and G. M. Acland, 2010 Exonic SINE insertion in STK38L causes canine early retinal degeneration (erd). Genomics 96: 362–368.

Hancks, D. C., and H. H. Kazazian, 2016 Roles for retrotransposon insertions in human disease. Mobile DNA 7: 9.

Ivancevic, A. M., R. D. Kortschak, T. Bertozzi, and D. L. Adelson, 2016 LINEs between Species: Evolutionary Dynamics of LINE-1 Retrotransposons across the Eukaryotic Tree of Life. Genome Biol Evol 8: 3301–3322.

Kirkness, E. F., V. Bafna, A. L. Halpern, S. Levy, K. Remington et al., 2003 The Dog Genome: Survey Sequencing and Comparative Analysis. Science 301: 1898–1903.

Lev-Maor, G., O. Ram, E. Kim, N. Sela, A. Goren et al., 2008 Intronic Alus Influence Alternative Splicing. PLOS Genetics 4: e1000204.

Li, H., and R. Durbin, 2009 Fast and accurate short read alignment with Burrows–Wheeler transform. Bioinformatics 25: 1754–1760.

Li, Z., M. K. Mulligan, X. Wang, M. F. Miles, L. Lu et al., 2010 A Transposon in Comt Generates mRNA Variants and Causes Widespread Expression and Behavioral Differences among Mice. PLOS ONE 5: e12181.

Lin, L., J. Faraco, R. Li, H. Kadotani, W. Rogers et al., 1999 The Sleep Disorder Canine Narcolepsy Is Caused by a Mutation in the Hypocretin (Orexin) Receptor 2 Gene. Cell 98: 365–376.

Lindblad-Toh, K., C. M. Wade, T. S. Mikkelsen, E. K. Karlsson, D. B. Jaffe et al., 2005 Genome sequence, comparative analysis and haplotype structure of the domestic dog. Nature 438: 803–819.

Marchant, T. W., E. J. Johnson, L. McTeir, C. I. Johnson, A. Gow et al., 2017 Canine Brachycephaly Is Associated with a Retrotransposon-Mediated Missplicing of SMOC2. Current Biology 27: 1573–1584.e6.

Minnick, M. F., L. C. Stillwell, J. M. Heineman, and G. L. Stiegler, 1992 A highly repetitive DNA sequence possibly unique to canids. Gene 110: 235–238.

Murphy, S. C., J. M. Evans, K. L. Tsai, and L. A. Clark, 2018 Length variations within the Merle retrotransposon of canine PMEL: correlating genotype with phenotype. Mobile DNA 9: 26.

Okutman, O., J. Muller, Y. Baert, M. Serdarogullari, M. Gultomruk et al., 2015 Exome sequencing reveals a nonsense mutation in TEX15 causing spermatogenic failure in a Turkish family. Hum. Mol. Genet. 24: 5581–5588.

Ostrander, E. A., F. Galibert, and D. F. Patterson, 2000 Canine genetics comes of age. Trends in Genetics 16: 117–124.

Pelé, M., L. Tiret, J.-L. Kessler, S. Blot, and J.-J. Panthier, 2005 SINE exonic insertion in the PTPLA gene leads to multiple splicing defects and segregates with the autosomal recessive centronuclear myopathy in dogs. Human Molecular Genetics 14: 1417–1427.

Ponicsan, S. L., J. F. Kugel, and J. A. Goodrich, 2010 Genomic gems: SINE RNAs regulate mRNA production. Current Opinion in Genetics & Development 20: 149–155.

Powell, J. A., J. Allen, and N. B. Sutter, 2010 DOG-SPOT database for comprehensive management of dog genetic research data. Source Code Biol Med 5: 10.

Ramkumar, N., B. M. Harvey, J. D. Lee, H. L. Alcorn, N. F. Silva-Gagliardi et al., 2015 Protein O-Glucosyltransferase 1 (POGLUT1) Promotes Mouse Gastrulation through Modification of the Apical Polarity Protein CRUMBS2 (A. Sutherland, Ed.). PLoS Genet 11: e1005551.

Silva, P., A. Justicia, A. Regueiro, S. Fariña, J. M. Couselo et al., 2017 Autosomal recessive agammaglobulinemia due to defect in μ heavy chain caused by a novel mutation in the IGHM gene. Genes Immun 18: 197–199.

Smith, B. F., Y. Yue, P. R. Woods, J. N. Kornegay, J.-H. Shin et al., 2011 An intronic LINE-1 element insertion in the dystrophin gene aborts dystrophin expression and results in Duchenne-like muscular dystrophy in the corgi breed. Lab Invest 91: 216–231.

Steenbeek, F. G. van, M. K. Hytönen, P. a. J. Leegwater, and H. Lohi, 2016 The canine era: the rise of a biomedical model. Animal Genetics 47: 519–527.

Sultana, T., D. van Essen, O. Siol, M. Bailly-Bechet, C. Philippe et al., 2019 The Landscape of L1 Retrotransposons in the Human Genome Is Shaped by Pre-insertion Sequence Biases and Post-insertion Selection. Molecular Cell 74: 555–570.e7.

Takeuchi, A., K. Iida, T. Tsubota, M. Hosokawa, M. Denawa et al., 2018 Loss of Sfpq Causes Long-Gene Transcriptopathy in the Brain. Cell Reports 23: 1326–1341.

Taskesen, M., G. B. Collin, A. V. Evsikov, A. Güzel, R. K. Özgül et al., 2012 Novel Alu retrotransposon insertion leading to Alstrom syndrome. Human Genetics; Heidelberg 131: 407–13.

Timmers, C., T. Taniguchi, J. Hejna, C. Reifsteck, L. Lucas et al., 2001 Positional Cloning of a Novel Fanconi Anemia Gene, FANCD2. Molecular Cell 7: 241–248.

Tsirigos, A., and I. Rigoutsos, 2009 Alu and B1 Repeats Have Been Selectively Retained in the Upstream and Intronic Regions of Genes of Specific Functional Classes (G. D. Stormo, Ed.). PLoS Comput Biol 5: e1000610.

Wang, W., and E. F. Kirkness, 2005 Short interspersed elements (SINEs) are a major source of canine genomic diversity. Genome Res. 15: 1798–1808.

Wiedmer, M., A. Oevermann, S. E. Borer-Germann, D. Gorgas, G. D. Shelton et al., 2016 A *RAB3GAP1* SINE Insertion in Alaskan Huskies with Polyneuropathy, Ocular Abnormalities, and Neuronal Vacuolation (POANV) Resembling Human Warburg Micro Syndrome 1 (WARBM1). G3 6: 255–262.

Yang, J.-J., P.-J. Liao, C.-C. Su, and S.-Y. Li, 2005 Expression patterns of connexin 29 (GJE1) in mouse and rat cochlea. Biochemical and Biophysical Research Communications 338: 723–728.

Zeng, R., F. H. G. Farias, G. S. Johnson, S. D. McKay, R. D. Schnabel et al., 2011 A Truncated Retrotransposon Disrupts the GRM1 Coding Sequence in Coton de Tulear Dogs with Bandera’s Neonatal Ataxia: Canine GRM1 Mutation Causes Neonatal Ataxia. Journal of Veterinary Internal Medicine 25: 267–272.

Zhang, Y., M. T. Romanish, and D. L. Mager, 2011 Distributions of Transposable Elements Reveal Hazardous Zones in Mammalian Introns (I. Rigoutsos, Ed.). PLoS Comput Biol 7: e1002046.

