## Supplement for "Polymorphic SINEC_Cf Retrotransposons in the Genome of the Dog (*Canis familiaris*)"

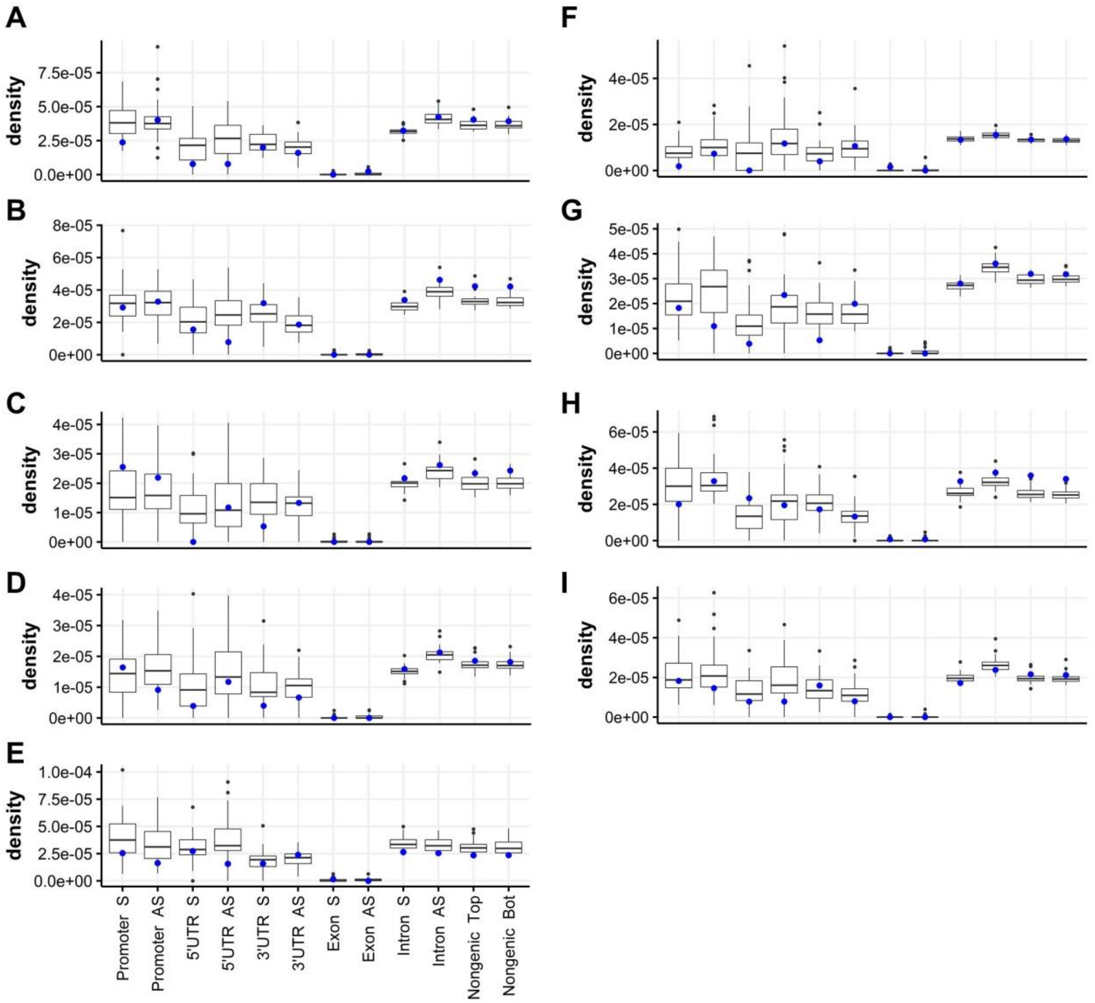


**Figure S1** SINE density varies by strand and part of gene. Each boxplot shows the SINE density (the number of reference genome annotated copies per ungapped bp) across the dog's 39 chromosomes, with chromosome X specially indicated in blue. Gene regions followed by "S" show the density of sense strand SINEs while "AS" shows the antisense density. For the nongenic (intergenic) category the single strands are top and bottom. The order of gene regions is the same in all panels. (A) SINEC_a1, (B) SINEC_a2, (C) SINEC_b1, (D) SINEC_b2, (E) MIR, (F) SINEC_c1, (G) SINEC_c2, (H) SINEC_Cf2, (I) SINEC_Cf3.

**Table S1** Oligonucleotides used for library construction and PCR-validation of randomly selected polymorphic SINEs.

**Table S2** Polymorphic SINEs present in at least one library and absent from the dog CanFam 3.1 reference genome.

**Table S3** Validated polymorphic SINEs.

**Table S4** Putative polymorphic SINEs in coding exons.
